# Patterned ELR-Gelatin Hydrogels Enable Rapid Endothelial Monolayer Formation via Bioactive Matrix Chemistry and Surface Topography

**DOI:** 10.64898/2026.03.22.713452

**Authors:** Jagoda Litowczenko, Yannick Richter, Martyna Michalska, Piotr Paczos, Karine Tadevosyan, Daniel Uribe, Jose Carlos Rodriguez-Cabello, Ioannis Papakonstantinou, Angel Raya

## Abstract

The endothelialization of organ-on-chip platforms and vascular implants is often limited by slow cell attachment and unstable monolayer formation. This work presents a scalable workflow that imprints micro- and nano-gratings into elastin-like recombinamer (ELR)-based hydrogels, enabling rapid endothelial cell capture and accelerating monolayer formation within 14 days. Three gelatin-ELR formulations are engineered, with ¹H-NMR confirming incorporation of sequences designed to modulate bioactivity (ELR1: inert; ELR2: uPA-responsive; ELR3: RGD-adhesive). ELR incorporation generates fibrillar microstructures and enhances mechanical performance, yielding elastic-dominant networks suitable for high-fidelity pattern transfer and stable culture. Using this library, the combined effects of ELR bioactivity and groove geometry on human iPSC-derived endothelial cells (iPSC-ECs) are systematically evaluated. In a 15-minute attachment assay, patterned ELR composites markedly improve cell retention compared to gelatin, with ELR2 on ∼350 nm and ∼4 µm grooves performing best, consistent with controlled, cell-mediated interfacial remodeling. This early advantage persists, as ELR2 and ELR3 hydrogels support rapid alignment and reach confluence by day 14, whereas gelatin remains sub-confluent. Cytoskeletal analysis confirms F-actin alignment. By combining enhanced early capture with protease-regulated remodeling, ELR2 identifies a favorable design window. These results establish a materials design framework linking programmable ELR chemistry with surface topography to engineer endothelial interfaces, providing a versatile platform for vascular biomaterials and microphysiological systems.

## INTRODUCTION

A confluent, quiescent endothelium is essential for controlling thrombosis, inflammation, and barrier function at blood–material interfaces. However, technologies such as organ-on-chip platforms and vascular implants (e.g., small-diameter grafts) are often limited by slow and unstable endothelialization. This is partly because the durable polymers most commonly used compromise endothelial attachment, long-term patency, and provide neither the biochemical nor topographical cues for stable monolayer formation [1], [18].

Surface topography is a potent physical cue: sub-micron to micron-scale parallel grooves can direct protrusion dynamics, focal-adhesion maturation, and cytoskeletal organization, promoting endothelial elongation, alignment, directed migration, and retention—even under shear [2]–[4], [9], [12]–[14], [19]. Yet, most parametric studies on topographical guidance have been conducted on rigid materials or PDMS leaving a gap in our understanding of how identical geometries influence cells on soft, protein-mimetic hydrogels that better emulate the native basement-membrane mechanics [19], [20].

To address this, we turn to elastin-like recombinamers (ELRs), which provide a sequence-defined route to soft matrices with modular bioactivity (e.g., RGD, protease-responsive motifs), mild processing conditions, optical clarity, and tunable elasticity [5]–[7], [21]. As opposed to topographies typically studied on static surfaces, this enables us to investigate them within degradable, cell-remodeled hydrogels. Concurrently, induced pluripotent stem cell–derived endothelial cells (iPSC-ECs) offer a renewable human cell source, but their sensitivity to the seeding microenvironment necessitates strategies that enhance rapid capture and accelerate time-to-confluence through stable early integrin anchorage [8], [11], [15], [16], [22].

Despite significant progress in hydrogel engineering, no existing platform combines optical clarity, rapid imprintability and compatibility with curved, lumen-like geometries—three prerequisites for generating pre-patterned endothelia capable of withstanding physiological shear stress in organ-on-chip systems and small-diameter vascular grafts. Addressing this unmet technological need requires materials that couple minute-scale endothelial capture with long-term monolayer stability under flow which we aim to engineer.

Here, we design an optically clear, rapidly imprintable gelatin–ELR composite hydrogel. By varying the ELR sequence (ELR1–3: ELR1, a non-bioactive baseline; ELR2, a protease-responsive variant specifically degradable by uPA; and ELR3, an RGD-containing adhesive variant), we tune both mechanics and bioactivity relative to a gelatin control, which is inherently cell-adhesive but lacks defined, tunable adhesion and remodeling cues. We then combine this chemistry with fast soft-lithographic imprinting to create a library of patterned hydrogels featuring nano- (N1/N2) and micro-scale (M1–M4) gratings of low aspect ratios (<1), alongside flat controls (F). This design allows us to systematically deconvolute the effects of biochemical composition (ELR sequence) and topography (grating spacing and depth) within a single, culture-compatible system. We selected two representative length scales that correspond to well-established endothelial sensing regimes. Micron-scale gratings (4–8 µm width; 0.5–1 µm height) fall within the dimensions repeatedly shown to guide endothelial elongation, junction alignment and collective migration on rigid and PDMS substrates [3,13], and include recent demonstrations of distinct guidance mechanisms in single cells versus monolayers on similar microstructured surfaces [9]. In contrast, the ∼350 nm width regime probes the sub-micron domain associated with integrin clustering, filopodial probing and nascent adhesion formation, which initiate contact-guidance responses on nanopatterned substrates [3]. Because most evidence for both scales originates from stiff materials, we used shallow aspect ratios (<1) to ensure reliable replication and optical clarity in soft ELR–gelatin hydrogels. This micro/nano subset therefore provides a biologically meaningful and fabrication-robust window for isolating geometric effects in compliant protein matrices.

Using this framework, we map how chemistry and geometry regulate: (i) initial adhesion (minutes), (ii) early contact guidance, and (iii) monolayer expansion over two weeks, and we established a deployable chemistry-geometry palette for rapid, stable patterned endothelialization. Finally, by optimizing the viscoelastic properties of the gelatin–ELR blend and using thin, flexible intermediate stamps, make this imprinting strategy inherently compatible with replication on gently curved surfaces and within tubular molds, extending this approach to the lumen-facing geometries relevant to vascular models and devices [2]–[4], [5]–[7], [8], [9], [11]–[17], [19], [23], [24].

Overall, we provide an experimentally grounded map of how ELR content enhances gelatin hydrogels to form rapid, stable endothelial monolayers, advancing strategies for vascular tissue engineering [2]–[4], [5]–[7], [8], [9], [11]–[16], and align with broader evidence that micro/nano-patterns regulate cellular behavior across lineages [17], [24]. Importantly, rather than focusing on a single mechanism, this work aims to establish a design framework for endothelial interface engineering, linking matrix bioactivity, mechanics, and surface geometry.

## MATERIALS AND METHODS

### Development of patterned ELR-based hydrogels

#### (i) Fabrication of nano/micro-patterned silicon masters

Nanogratings and microgratings were defined on Si wafers by Lloyd’s-mirror laser interference lithography (LIL) and direct laser writing (Heidelberg DWL 66+ laser writer lithography system), respectively. Wafers were solvent-cleaned, O₂-plasma activated (100 W, 60 s, O_2_ 7 sccm), hexamethyldisilazane (HMDS)-primed, spin-coated with a photoresist (ma-N 405 at 300 nm thickness for LIL and S1818 at 1.8 µm for DWL), soft-baked, then exposed via LIL to achieve 700 nm pitch or via DWL for 8 and 16 µm pitches. After development, patterns were transferred into silicon by Bosch process in Inductively Coupled Plasma-Reactive Ion Etching (ICP-RIE) to the target heights H of ∼1.0 and ∼0.5 µm [30].

##### Pattern encoding and nomenclature

To ensure consistent terminology across all experiments, we refer to each topographical pattern using a unified abbreviated code in the main manuscript and figures. The full structural parameters and their corresponding codes are: F (Flat), N1 (N350H1), N2 (N350H0.5), M1 (M4H1), M2 (M8H1), M3 (M4H0.5), and M4 (M8H0.5).

Codes denote the groove width (N = 350 nm; M = 4–8 µm) and relief height (H1 = 1 µm; H0.5 = 0.5 µm). The gratings have duty cycle of 0.5, thus grove and grating widths are the same corresponding to pitch of 700 nm, 8 and 16 µm. Full mapping is provided in ***Supplementary Table S1***, which includes groove width and height parameters for clarity.

#### (ii) Engineering of ELR-based hydrogels

We engineered imprintable composite hydrogels by blending gelatin with elastin-like recombinamers selected to decouple basal mechanics from bioactivity. Three ELRs were used throughout: VKV24 (no specific adhesion or protease-sensitive motifs; referred to hereafter as ELR1), DRIR (contains the slowly urokinase-type plasminogen activator (uPA)-degradable DRIR sequence; ELR2), and HRGD6 (presents RGD adhesion motifs; ELR3). Since endothelial cells orchestrate local fibrinolysis and pericellular matrix remodeling through the uPA/uPAR system, which plays a key role in VEGF-driven angiogenesis and endothelial migration [29], embedding uPA-degradable sequences within ELRs provides a physiologically relevant mechanism for cell-mediated remodeling of the coating. This, in turn, is expected to facilitate endothelial spreading, migration, and stable monolayer formation under vascular-like conditions. These sequences and their biophysical rationale follow prior ELR design frameworks and were produced and characterized as previously reported for VKV-, DRIR- and RGD-bearing ELRs [26], [27].

##### Stock solutions

Type A porcine gelatin (Sigma G1890) was dissolved in DMEM, sterile-filtered (0.22 µm). ELRs were dissolved at 50 mg mL⁻¹ in PBS (0.22 µm filtered). Phases were combined 0.75 (gelatin/DMEM) : 0.25 (ELR/PBS) and cycled on ice/RT/30 °C until homogenous and slightly opalescent.

##### Crosslinking

Microbial transglutaminase (mTG) was added immediately before casting (manufacturer’s activity; details in Supplementary Methods), forming ε-(γ-glutamyl)-lysine bonds between gelatin and ELR lysines.

#### (iii) Pattern transfer into ELR hydrogels

To fabricate patterned ELR hydrogels, three lithographic steps were followed: (iii-i) transfer from masters to Intermediate Polymer Stamps (IPS) via nanoimprint lithography (NIL) to preserve the master longevity, (iii-ii) transfer from IPS to poly(urethane acrylate) (PUA) for the process scale up (number of IPS was limited; please note this step can be omitted if the number of IPS is sufficient), and (iii-iii) transfer from PUA to ELR composite by stamping.

#### (iii-i) NIL (master → IPS)

Briefly, prior to NIL, masters were silanized in trichlorooctadecylsilane for 60 min (1% v/v toluene solution) and post-annealed (120 °C, 30 min) to facilitate demolding [28]. Pattern dimensions were verified by scanning electron microscopy (SEM, Zeiss) and optical profilometry (Keyence VHX-series microscope) before replication. The NIL process was performed on EITRE 3 (Obducat). The IPS (Obducat) stamping process was operated at 145 °C and 30 bar for 20 s.

#### (iii-ii) Soft-lithographic casting of PUA daughter stamps (IPS → PUA)

PUA resin was compounded according to our previous protocols (ACSNano 2020,14,12091−12100). Briefly, diacrylate prepolymer (Ebecryl 284, Allnex) was blended with 30 wt% trimethylolpropane-ethoxylate triacrylate (Sigma 409073) as reactive diluent; photoinitiators Irgacure 184 (BASF 30472119) and 2-hydroxy-2-methylpropiophenone (Sigma 405655) were each added at 1.5 wt% relative to total resin; TEGO® Rad 2200N (1 wt%) improved wetting/release. The mixture was vortexed, degassed and stored cold, protected from light. A togetherilm was drop-cast on IPS, overlaid with a clean glass support to define a uniform, thin stamp, and UV-cured for ∼20 s at 365 nm (high intensity), followed by a 10 h low-intensity post-cure. Stamps were gently demolded to yield flexible PUA negatives suitable for repeated use. Before cell work, stamps were rinsed in isopropanol, air-dried, sterilized in 70% ethanol and UV-irradiated.

#### (iii-iii) Soft-lithographic stamping of ELR-based hydrogels (PUA → ELR hydrogel)

Freshly prepared ELR–gelatin (with mTG) blends were dispensed into 48-well tissue-culture plates at 0.50 mL per well using a positive-displacement pipette to ensure volumetric accuracy and minimize bubble entrainment. After casting, plates were left at room temperature of 21 °C (RT) for 3 min to allow incipient gelation (pre-set), providing sufficient viscoelasticity for imprinting without loss of conformity.

For topography transfer, a thin PUA stamp carrying the line pattern was gently placed pattern-side down onto each pre-set hydrogel surface, ensuring full areal contact and avoiding lateral sliding. The hydrogel–stamp assemblies were maintained at RT for 10 min, then transferred to 4 °C for a further 10 min to stabilize the imprinted interface while the mTG reaction progressed. Plates were subsequently removed to RT and kept undisturbed for an additional ∼40 min to allow completion of crosslinking. Stamps were peeled, gels rinsed in PBS and equilibrated in medium prior to seeding. All steps were performed under sterile conditions; stamps were handled with sterile forceps and lowered vertically to avoid trapping air. Minimal pressure was applied only to ensure conformal contact; no external load was used.

### Hydrogel characterization

#### Oscillatory rheology

Small-amplitude oscillatory rheology was performed on a stress-controlled rheometer (Anton Paar MCR-302) using a 25 mm serrated parallel-plate geometry with a gap of 1.0 mm. Samples were loaded at 25 °C, trimmed, and equilibrated for 5 min. Strain-sweep tests (0.01–100%) at 1 rad s⁻¹ identified the linear viscoelastic regime (LVR). Frequency sweeps (0.1–100 rad s⁻¹) were then run at γ = 0.5% (within LVR). Temperature was controlled with a Peltier stage (±0.1 °C). Storage (G′), loss (G″) moduli, and complex viscosity (η*) were exported as mean ± SD across n = 3 hydrogels per formulation.

#### Uniaxial tensile testing

Dog-bone-like strips (width 5 mm, gauge length 15 mm, thickness 1.5–2 mm) were cut from fully cross-linked gels and tested in tension on a Zwick/Roell Z100 with a 5 kN load cell (class 1 accuracy to 0.2% Fnom). Custom flat grips used 61 HRC steel inserts coated with 95 ShA polyurethane to prevent slippage/necking of soft gels. Tests were run at 60 mm min⁻¹ from a 0.1 N pre-load to failure. True stress–strain curves were computed by engineering-to-true conversion assuming incompressibility (Poisson ∼0.5) within the large-strain regime. We report ultimate tensile strength (UTS), strain at break, Young’s modulus (tangent at 5–10% strain), and stiffness (N mm⁻¹); n = 4 per material.

#### ¹H NMR of composites

Proton spectra (¹H NMR) of ELR components and Gelatin-ELR composites were acquired on a Bruker AVANCE III 400 MHz at 298 K using standard single-pulse acquisition with water presaturation. Chemical shifts are referenced to TSP (0.00 ppm) in D₂O. Spectra were phase- and baseline-corrected in MestReNova, then overlaid after intensity normalization. Representative fingerprints differentiating VKV24 (ELR1) and HRGD6 (ELR3) are shown in Fig. 2L. For ELR2, the user provided a re-exported ¹H spectrum after an initial file that opened as ¹³C (100–200 ppm range); the final overlay is included in Fig. 2L. Full peak lists are in Supplementary Data.

### Human iPSC culture

All work with human iPSC lines was conducted under approval from the Spanish competent authorities (Commission on Guarantees concerning the Donation and Use of Human Tissues and Cells, Instituto de Salud Carlos III). The FiPS Ctrl1-mR5F-6 line (registered at the Spanish National Stem Cell Bank) was maintained on Matrigel®-coated cultureware (Corning) in mTeSR1 medium (STEMCELL Technologies). Medium was renewed daily except on the first day after passaging. For routine expansion, colonies were split 1:4–1:6 by brief incubation with 0.5 mM EDTA (Invitrogen) at 37 °C (∼2 min) and re-plated as small aggregates onto fresh Matrigel for continued maintenance.

### Differentiation of endothelial cells

Endothelial differentiation was adapted from established lateral-mesoderm protocols [8]. hiPSCs on Matrigel were dissociated to single cells with Accutase (37 °C, ∼8 min) and seeded onto Matrigel-coated 12-well plates at 1.5 × 10^5^ cells/well in mTeSR1 with 10 µM Y-27632 (ROCK inhibitor, Sigma). After 24 h, medium was switched to EC priming medium (N2B27 composed of DMEM/F-12:Neurobasal 1:1 with N2 and B27 supplements) containing 8 µM CHIR99021 (GSK3β inhibitor; Sigma, SML1046) and 25 ng ml⁻¹ BMP4 (R&D Systems) for 3 days. Cultures were then transferred to EC induction medium (StemPro-34 SFM; Thermo Fisher) supplemented with 200 ng ml⁻¹ VEGF (PeproTech) and 2 µM forskolin (Sigma-Aldrich) for a further 3 days.

### Expansion and purification of endothelial cells

On day 6, cells were re-plated onto 0.1% gelatin-coated dishes and expanded in EGM-2 (Lonza) supplemented with a YAC cocktail (3 µM CHIR99021, 10 µM Y-27632, and 10 µM SB431542) for ∼4 days. Purification was performed by sequential differential detachment with TrypLE (Thermo Fisher): after two PBS rinses, cultures received TrypLE at room temperature until non-EC contaminants detached with gentle tapping and were removed; remaining ECs were then exposed to TrypLE at 37 °C (∼5 min), collected, and re-plated on 0.1% gelatin. iPSC-ECs at passages 2–3 were used for all experiments.

### Immunostaining analysis

Hydrogel constructs (or control coverslips) were fixed in 4% paraformaldehyde (PFA; Sigma) for 15–30 min (RT or 4 °C), rinsed in TBS (3 × 10–30 min) and permeabilized/blocked in TBS with 0.1% Triton X-100 and 6% donkey serum (Chemicon). Primary antibodies were diluted in blocking buffer and incubated at 4 °C with gentle agitation (overnight to 72 h depending on target). After 4 TBS washes, fluorescent secondary antibodies were applied for 2 h at RT (light-protected) or overnight at 4 °C, followed by DAPI nuclear counterstaining. F-actin was labeled with fluorescent phalloidin. Imaging was performed on a Zeiss LSM980 Airyscan2; image processing used FIJI/ImageJ and, where indicated, Imaris (Microscopy Image Analysis Software, Oxford Instruments). Primary and secondary antibodies used are listed in **Table S2.**

### Quantitative real-time polymerase chain reaction (qRT-PCR)

Total RNA was extracted from iPSC-ECs using Maxwell® RSC simplyRNA Cells kits on a Maxwell RSC instrument (Promega). cDNA was synthesized from 1 µg RNA using Transcriptor First Strand cDNA kit (Roche). qPCR was run on a LightCycler® 480 (Roche) with GAPDH as housekeeping control. RNA from primary HUVECs served as a positive endothelial control, and human dermal fibroblast RNA as a negative control. Primer sequences are listed in **Table S2**.

### Live/dead assay

Live/dead working solution was prepared in PBS by mixing 50 µl propidium iodide (PI; 2 mg ml⁻¹ stock in PBS; Sigma-Aldrich) and 8 µl fluorescein diacetate (FDA; 5 mg ml⁻¹ stock in acetone; Sigma-Aldrich). Constructs were rinsed with PBS, incubated with the staining mixture for up to 5 min at RT in the dark, washed with PBS, and imaged immediately in phenol red free DMEM by confocal microscopy.

### Hydrogels topography, cell morphology, and spreading analysis

Bright-field images (attachment and coverage assays) were acquired on a digital microscope (Keyence VHX-series) using identical exposure settings across conditions. iPSC-EC surface coverage (% area) was quantified in FIJI using background subtraction, auto-thresholding (Otsu), and binary area fraction within a fixed region of interest; four non-overlapping fields per condition were analyzed and averaged per well/biological replicate. Orientation was assessed from F-actin or phase-contrast images using OrientationJ, reporting angle histograms (0–180°) and rose plots; alignment indices were computed as described in Methods (binning and smoothing parameters as in Supplementary Information). Acquisition parameters (objective, pixel size, z-sampling) were held constant within each experiment.

### 15-minute attachment assay (retrieved-cell quantification)

iPSC-ECs were seeded at 150 000 cells/cm² in 48-well plates containing equilibrated hydrogels. After exactly 15 minutes, wells were gently rinsed once with 200 µL PBS to collect non-adherent (retrieved) cells. The PBS containing detached cells was transferred into microcentrifuge tubes, pelleted at 300 × g for 5 minutes, resuspended in 200 µL of medium, and counted using an automated cell counter. Retrieved-cell number was used as an inverse metric of adhesion strength (fewer retrieved cells = stronger immediate attachment). This assay quantifies early adhesion only.

### Quantification of cell orientation (OrientationJ → polar plots)

Based on phase-contrast or F-actin images, local orientation was computed in Fiji using the OrientationJ plugin (structure tensor). Angle histograms (0–180°) were exported and normalized to the sum of bin counts. For visualization, Python/Matplotlib generated polar bar charts with the zero-angle at the top and the axis displayed from −90° to +90°. Plot opacity/border width were standardized across conditions.

### Image processing and area-coverage calculation

Confocal/bright-field images were analyzed by an automated Python pipeline: 1. Preprocessing: grayscale import. 2. Adaptive thresholding: Gaussian method (blockSize = 11, C = 2) to segment cells under non-uniform illumination. 3. Morphology: closing then opening with a 5×5 rectangular kernel to fill intrusions and remove speckle. 5. Contour detection & hole filling: external contours were detected (cv2.RETR_EXTERNAL) and filled (thickness=FILLED). 6. Metric: area covered (%) = 100 × (cell-pixels)/(total pixels), where cell-pixels are non-zero entries of the filled mask. 7. Reporting: side-by-side panels (original/thresholded/filled) and per-image coverage were compiled into a multi-page PDF; a tab-delimited file (image name, % coverage) was saved for statistics. 8. Implementation used OpenCV, NumPy and Matplotlib; parameters were held constant within an experiment.

Area coverage (%) was analyzed per day by two-way ANOVA with factors Material (Gelatin, ELR1–3) and Pattern (F, N1/N2, M1–M4), including the interaction. Assumptions were evaluated on model residuals (Shapiro–Wilk; Brown–Forsythe). When needed, data were logit-transformed. Post-hoc contrasts used Šídák correction for (i) within-material comparisons (each pattern vs Flat) and (ii) within-pattern comparisons (each ELR vs Gelatin). As complementary kinetics metrics, we computed AUC(0–14 d) for %coverage and the time to 90–95% coverage (T90/T95) by linear interpolation between timepoints.

#### Orientation analysis (OrientationJ) and area coverage (Python)

OrientationJ (Fiji, structure tensor), normalization to the sum of bins, polar plots in Matplotlib with 0° at top and axis −90° to +90°. Coverage pipeline: grayscale import → adaptive Gaussian thresholding (blockSize = 11, C = 2) → morphological close/open (5×5 kernel) → external-contour fill → % area = non-zero / total × 100; panels and TSV exported; code uses OpenCV, NumPy, Matplotlib.

### Statistical analysis

Unless specified, data are mean ± s.d.; technical fields per well were averaged before inference. For 15-min retrievals (Fig. 6) and for coverage (Fig. 7), we applied two complementary approaches:

#### Primary (per-day) analysis

two-way ANOVA at each time point with factors Material (Gelatin, ELR1, ELR2, ELR3) and Pattern (F, N1, N2, M1–M4), Type III SS; residual diagnostics (Shapiro–Wilk; Brown–Forsythe); log₁₀ transform if needed. Post-hoc: Šídák for simple effects (pattern vs flat within material; ELR vs gelatin within pattern).

#### Global analysis (report in SI)

linear mixed-effects model with fixed effects Material, Pattern, Day (3/7/14) and all interactions; random intercept for replicate/well (fields nested), Satterthwaite df; post-hoc Tukey/Šídák. Provide partial η² (ANOVA) or conditional R² (LMM) as effect sizes and adjusted CIs.

All tests in GraphPad Prism 10 or R (lme4/emmeans); α = 0.05.

## RESULTS

### A scalable workflow for producing patterned ELR hydrogels

We established a scalable pipeline to generate line-patterned gelatin-ELR hydrogels. Using soft lithography, we replicated a library of micro- and nano-gratings from silicon masters onto intermediate polymer stamps (IPS), and finally into polyurethane acrylate (PUA) daughter stamps for reliable imprinting (**Figure 1**). This method faithfully transferred topographies across centimeter-scale areas, creating a library of patterns from nanometer (N1/N2) to micrometer (M1–M4) scales, alongside flat controls (F), suitable for high-resolution live-cell imaging.

**Figure 1.**
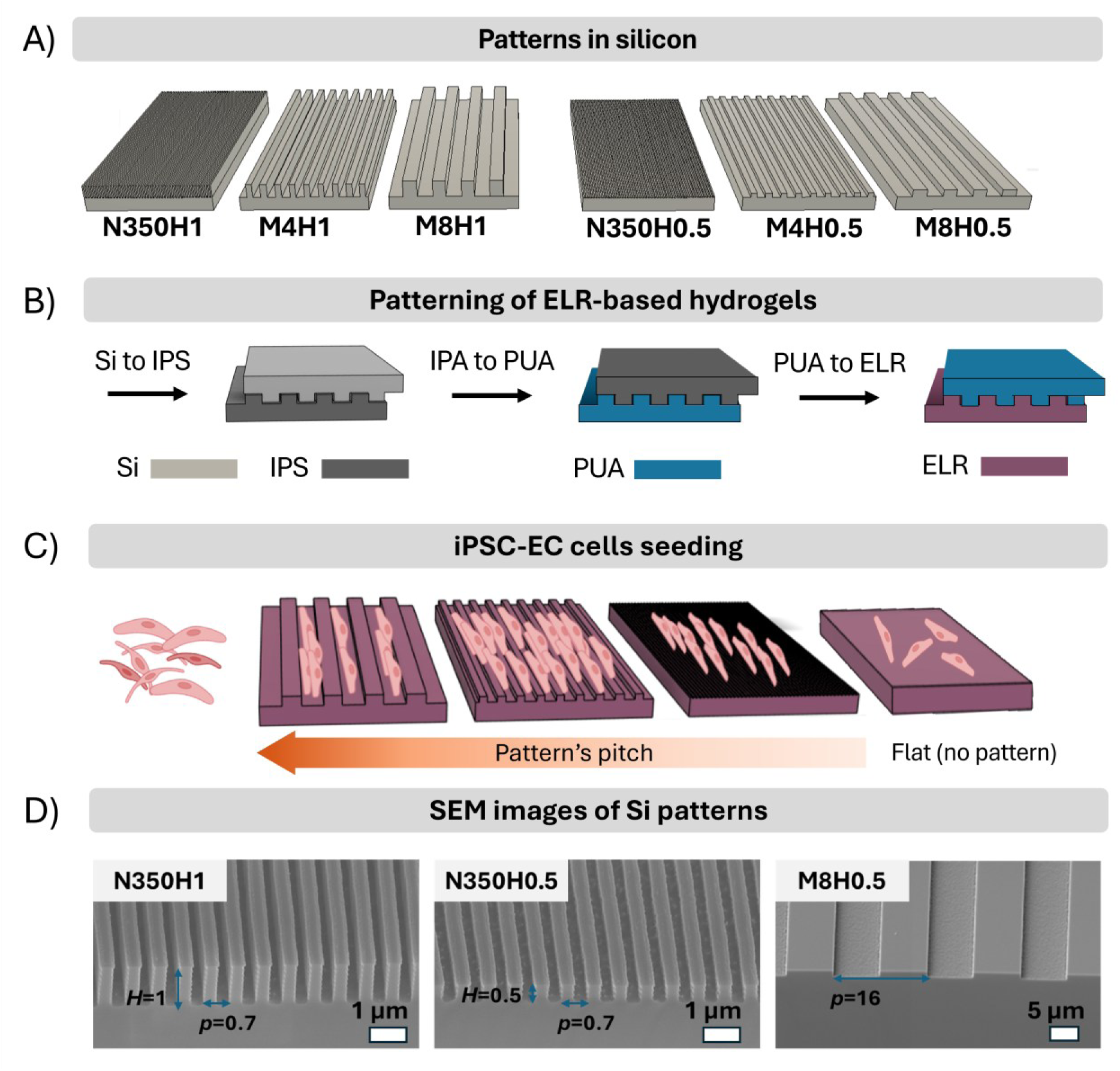
| Overview of the fabrication pipeline and experimental workflow. (A) Patterns in silicon: N350H1 (N1), M4H1 (M1), M8H1 (M2), N350H0.5 (N2), M4H0.5 (M3), and M8H0.5 (M4), corresponding to 350 nm, 4 µm, and 8 µm gratings with heights of ∼0.5 or 1 µm, fabricated by LIL/DWL and ICP-RIE. (B) Three-step pattern transfer workflow: Si → IPS (NIL), IPS → PUA, and PUA → ELR hydrogel. (C) Seeding of iPSC-derived endothelial cells (iPSC-ECs) onto patterned hydrogels, followed by readouts for immediate attachment, alignment, and coverage. (D) Representative patterned ELR hydrogels after imprinting and demolding. (E) SEM micrographs of silicon masters (N350H1, N350H0.5, M8H0.5). Scale bars as indicated.

Profilometry confirmed excellent pattern fidelity for micron-scale gratings, while nanoscale grooves (350 nm) showed reduced feature height, indicating partial amplitude loss during imprinting into the viscoelastic hydrogel. Pattern replication was robust across all material formulations **(Supplementary Fig. S1–S10).**

### ELR composites enhance mechanical and structural properties

We synthesized three gelatin-ELR hydrogels via enzymatic crosslinking with microbial transglutaminase, incorporating ELRs with distinct functionalities: VKV24 (ELR1, bioinert baseline), DRIR (ELR2, uPA protease responsive), and HRGD6 (ELR3, RGD-adhesive). ¹H-NMR confirmed successful ELR incorporation, indicated by increased signal at ∼0.9 ppm (valine isopropyl methyl groups) and distinct signatures of ELR2/3 at ∼1.7 ppm (isoleucine side chains) (**Fig. 2K-L**).

**Figure 2.**
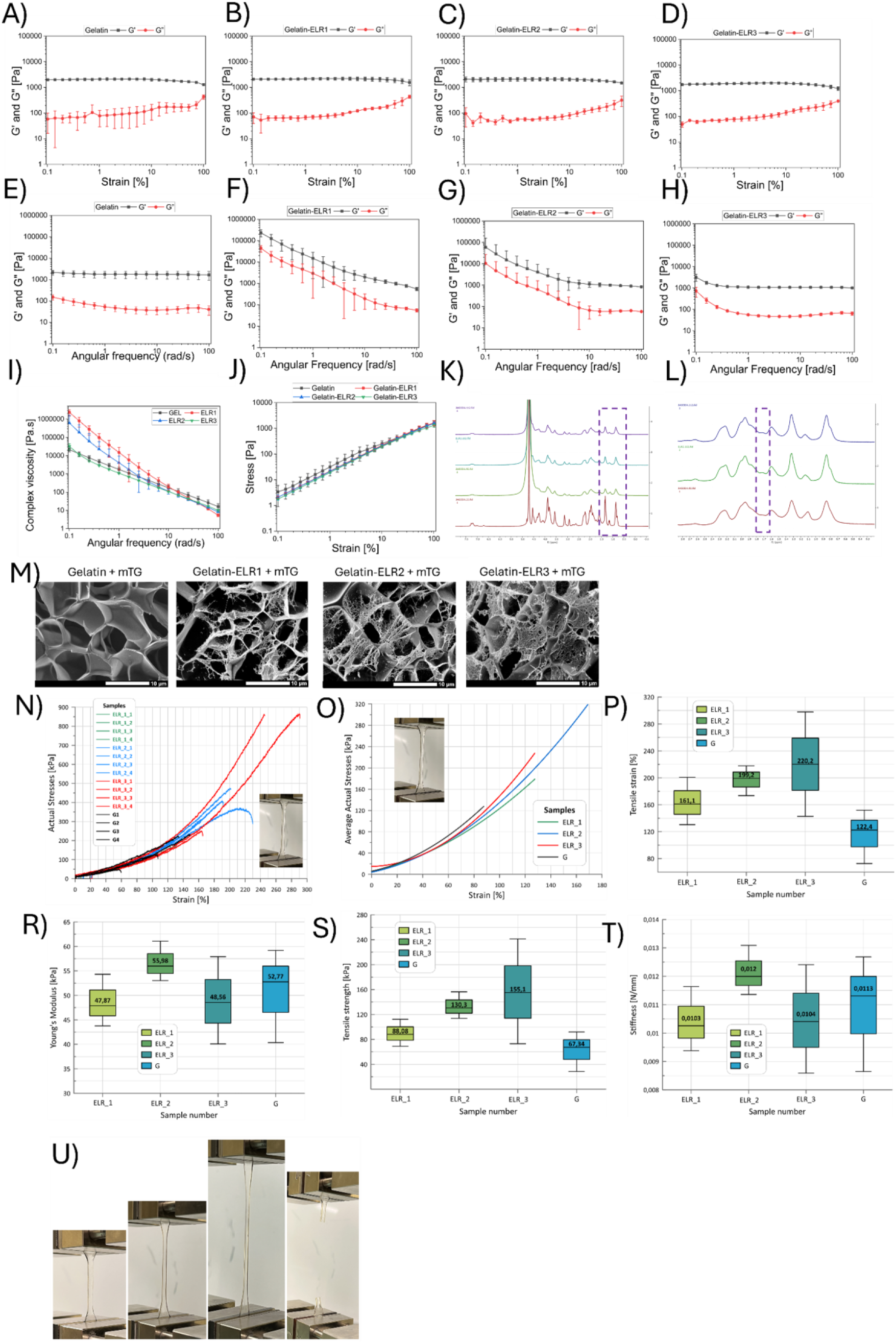
| Mechanical and molecular characterization of gelatin–ELR hydrogels. (A–H) Amplitude and frequency sweeps (γ = 0.5%, 0.1–100 rad s⁻¹) demonstrate solid-like viscoelastic behavior (G′ > G″) across all formulations. (I–J) Summary plots of storage modulus (G′), complex viscosity (η*), and stress response vs frequency (mean ± SD, n = 3). (K–L) ¹H-NMR overlays of gelatin and gelatin–ELR composites. Panel K highlights the ∼0.9 ppm region, where increased intensity in ELR1, ELR2 and ELR3 reflects valine isopropyl protons, confirming ELR incorporation. Panel L shows a zoom of the ∼1.7 ppm region, where ELR2 and ELR3 exhibit a more pronounced shoulder attributed to isoleucine methylene protons, consistent with their higher Ile content compared with ELR1 (VKVx24). These resonances collectively verify blending and allow molecular differentiation between ELR variants. (M) Cryo-SEM micrographs reveal a porous, fibrillar architecture in all hydrogels. (N–O) True stress–strain curves (individual and averaged) from uniaxial tension tests (Zwick Z100, 60 mm min⁻¹). (P–R) Box plots summarizing Young’s modulus, ultimate tensile strength, and stiffness (n = 4). ELR2 exhibits the most consistent modulus and stiffness, whereas ELR3 maximizes tensile strength. Scale bars as indicated.

Rheological characterization within the linear viscoelastic regime (γ = 1%) confirmed stable, gel-like behavior (G′ > G″) for all formulations (Fig. 2A-I). ELR incorporation enhanced the matrix resistance to deformation, with ELR2 and ELR3 composites sustaining higher stresses at matched strains than gelatin or ELR1 (Fig. 2E). Frequency sweeps and complex viscosity profiles were consistent with robust, shear-thinning viscoelastic networks (Fig. 2J).

Tensile testing revealed that ELR composites exhibited elastomeric, non-linear stress-strain curves, elongating beyond 100% strain before rupture (Fig. 2N–O, U). ELR3 achieved the highest ultimate tensile strength, albeit with wide variability, while ELR2 provided the most consistent and highest Young’s modulus, offering an favorable balance of stiffness and reproducibility (Fig. 2R–S). Cryo-SEM imaging revealed that ELR incorporation transformed the gelatin matrix into a more open, fibrous, and interconnected network, with ELR2 and ELR3 displaying finer fibrillar meshes consistent with their superior mechanical performance (**Fig. 2M**). Collectively, these data show that incorporating ELRs stabilizes the gelatin matrix against large-amplitude damage while preserving culture-relevant, elastic-dominant mechanics (Fig.2A–J) [18]-[21], [27].

### Generation and characterization of iPSC-derived endothelial cells

We differentiated iPSC into endothelial cells (iPSC-ECs) using a Wnt/BMP-guided protocol [8] (**Fig. 3A-G**). The resulting cells exhibited cobblestone morphology, formed continuous VE-cadherin-positive junctions (**Fig. 3H**), and expressed canonical endothelial markers (**Fig. 3I-J**): a junctional cadherin (CDH5, PECAM1/KDR) at levels comparable to HUVECs, confirming a stable endothelial phenotype suitable for downstream assays [8].

**Figure 3.**
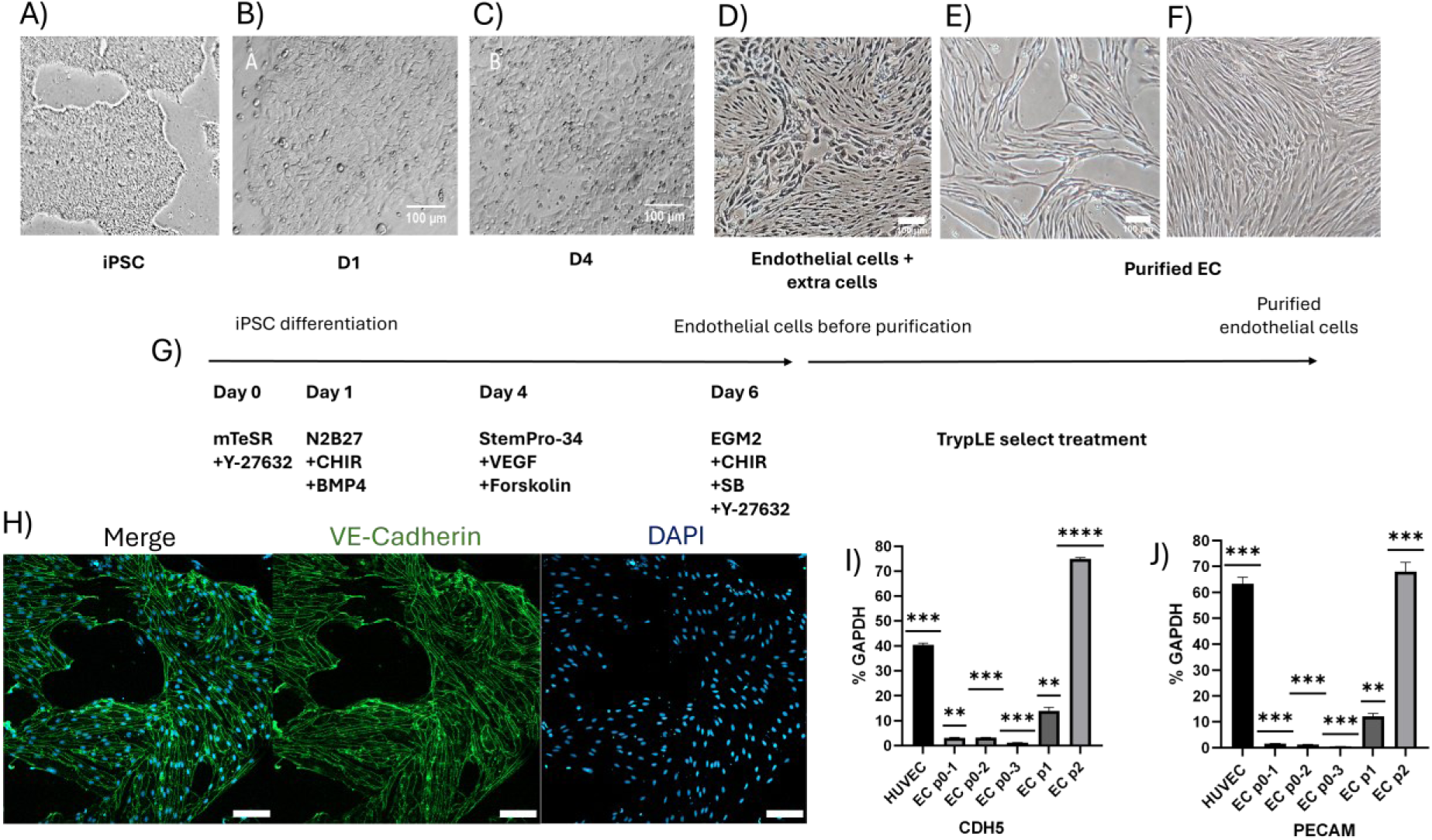
| Generation and characterization of human iPSC-derived endothelial cells (iPSC-ECs). (A–D) Bright-field images illustrating morphology at day 0, 1, 4, 6 of differentiation. (E) Post-selection/purification on day 6 and morphology (F) after expansion to day 12. (G) Schematic of the differentiation workflow adapted from established protocols. (H) Immunostaining at passage 2 showing junctional VE-cadherin (green); nuclei counterstained with DAPI (blue). Scale bar, 50 µm. (I) qRT-PCR analysis of endothelial marker CDH5 expression across passages 1–2; HUVEC serves as a positive control. Data are Mean ± SD (n = 3). (J) qRT-PCR analysis of endothelial marker PECAM1 expression across passages 1–2; HUVEC serves as a positive control. Data are Mean ± SD (n = 3).

### Combined effects of matrix chemistry and surface topography on immediate cell attachment

A critical 15-minute attachment assay revealed that both hydrogel chemistry and surface topography profoundly influenced initial cell adhesion. ELR-based surfaces retained significantly more cells than gelatin across all patterns (**Fig. 4A–E**). Quantitative analysis showed a several-fold reduction in retrieved (non-attached) cells on ELR substrates compared to gelatin (**Fig. 4B**). Patterns significantly modulated early cell retention. Among all geometries, the micron-scale M1–M2 grooves produced the lowest number of retrieved cells, indicating the strongest immediate attachment, followed by the sub-micron N1–N2 patterns, whereas Flat controls consistently showed weaker retention.

**Figure 4.**
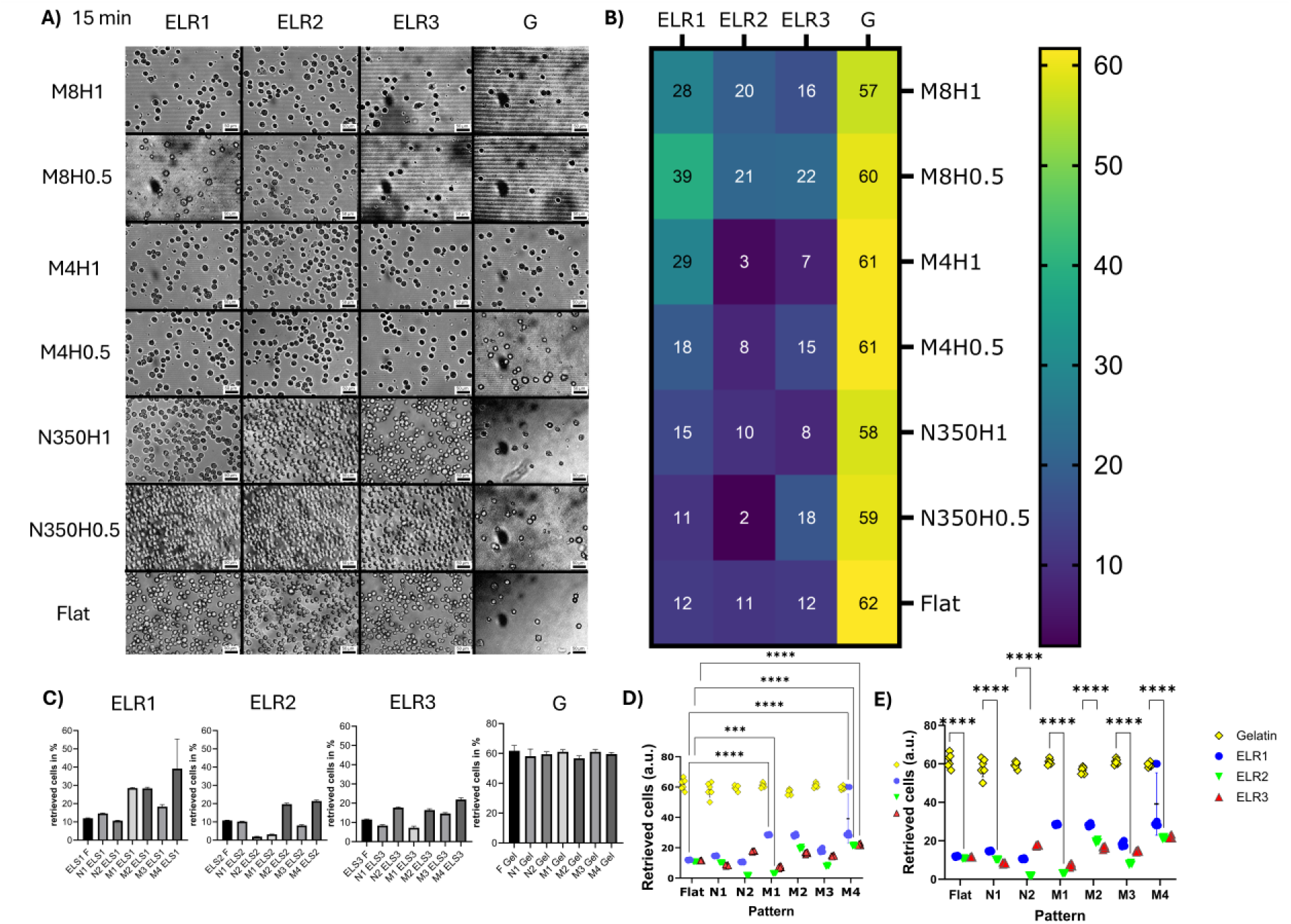
| Impact of topography and ELR formulation on immediate iPSC-EC attachment (15 min). (A) Representative bright-field micrographs 15 min after seeding, after removal of non-adherent cells by gentle washing. Rows show full pattern identifiers (M8H1, M8H0.5, M4H1, M4H0.5, N350H1, N350H0.5, Flat), columns show coating materials (ELR1–3, Gelatin). (B) Heat map of retrieved cells remaining in the supernatant (lower values = stronger immediate attachment). Tile numbers indicate mean values (a.u.). (C) Quantification of retrieved cells for each coating–pattern combination (mean ± SD; n = 6). (D) Comparison of retrieved cells across abbreviated pattern groups (F, N1–N2, M1–M4). (E) Comparison across coating materials (Gelatin, ELR1–3). Statistical analysis: two-way ANOVA with Šídák’s multiple-comparisons test (* P < 0.05, ** P < 0.01, *** P < 0.001, **** P < 0.0001). Note: Panels A–C display full pattern names; panels D–E use abbreviated codes (F, N1–N2, M1–M4) for readability. Mapping between full names and abbreviations is provided in Supplementary Table S1.

Across materials, ELR2 yielded the lowest retrieved-cell counts for all patterns, consistent with its uPA-responsiveness and enhanced pericellular remodeling capability. Importantly, comparison of flat conditions shows that all ELR formulations already enhance initial attachment relative to gelatin (Fig. 4B), establishing a chemistry-driven baseline. The further reduction in retrieved cells observed upon patterning—most pronounced for ELR2 on M1–M2 and N2 geometries—indicates that topographical guidance provides an additional, geometry-dependent contribution to early adhesion.

This chemistry-driven baseline was further modulated by topography. Specifically, the sub-micron N2 and micron-scale M1 grooves further enhanced attachment in combination with ELR chemistry. ELR2 (DRIR) performed exceptionally well on these patterns, yielding the lowest cell retrieval (**Fig. 4B-C**). Two-way ANOVA confirmed significant independent and interactive effects of both material and pattern (**Fig. 4D**). These results indicate that a protease-responsive, slowly remodeled interface (ELR2) particularly when combined with selected groove regimes, provides favorable conditions for nascent adhesion formation. The magnitude and speed of this effect are particularly relevant for perfused systems, where the first minutes after seeding determine long-term lumen coverage. Such rapid attachment under minimal shear-like washing steps indicates that these chemistries and groove regimes could markedly reduce cell loss during initial perfusion in chips and tubular grafts. Together, these observations support a model in which ELR chemistry defines the baseline adhesion capacity, while surface topography modulates the magnitude and spatial organization of early cell attachment.

### Rapid contact guidance and accelerated monolayer formation

Within hours of seeding, iPSC-ECs on patterned ELR hydrogels exhibited pronounced contact guidance. Sub-micron and micron groove patterns induced rapid axis alignment of iPSC-ECs, most prominently on M1–M2, followed by N1–N2, whereas Flat substrates produced isotropic spreading. Cells on micron-scale grooves (M1-M4) showed strong elongation and alignment parallel to the groove axis, while sub-micron patterns (N1/N2) induced a weaker, broader alignment. Flat controls were isotropic (**Fig. 5A**).

**Figure 5.**
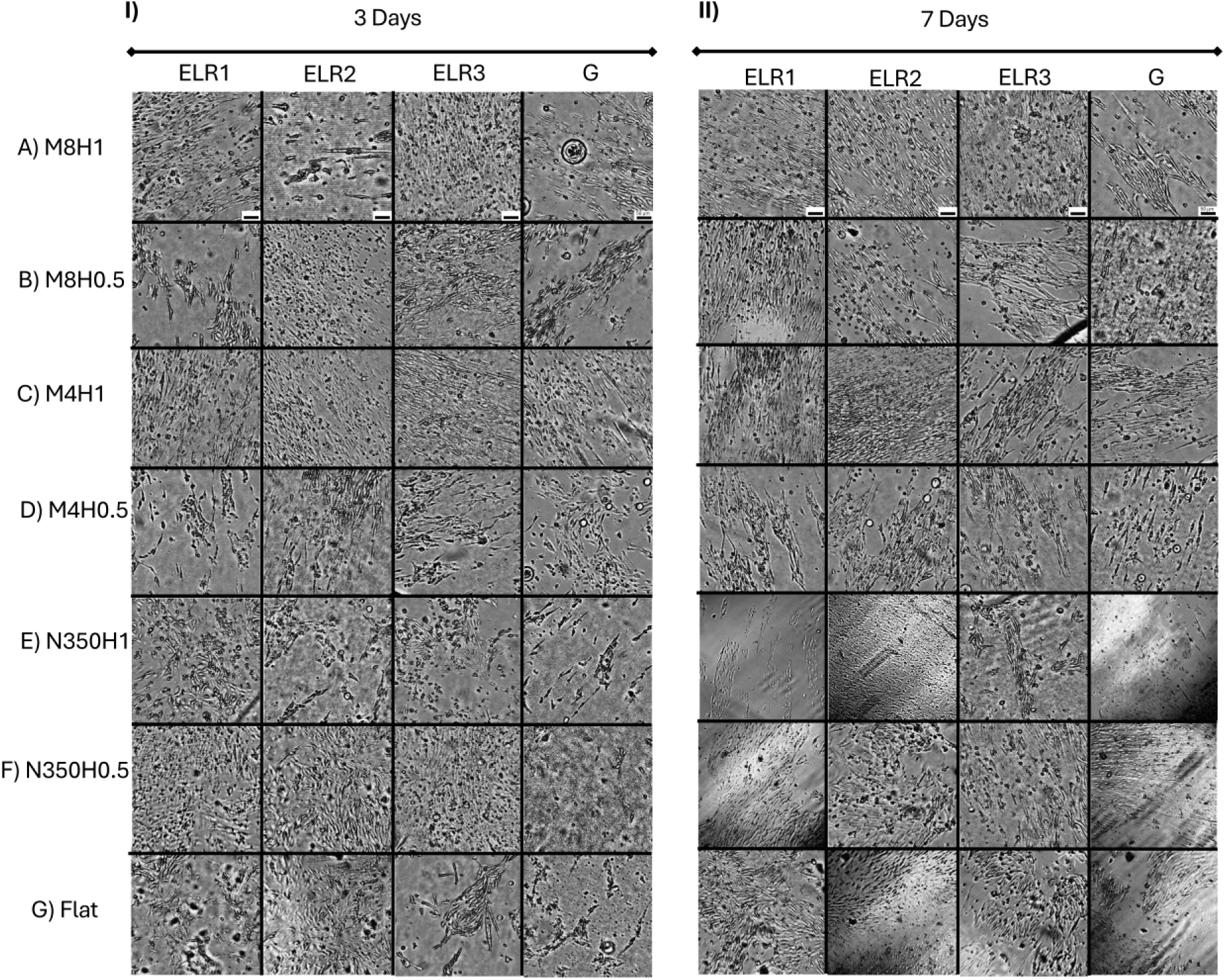
| Topography drives early orientation of iPSC-ECs on ELR hydrogels. (A) Representative phase-contrast images of iPSC-EC morphology and local alignment on patterned ELR hydrogels after 3 days of culture (same panel order as in Fig. 6). (B) Representative phase-contrast images of iPSC-EC morphology and local alignment on patterned ELR hydrogels after 7 days of culture. Scale bars as indicated. Pattern key: F (Flat), N1 = N350H1, N2 = N350H0.5, M1 = M4H1, M2 = M8H1, M3 = M4H0.5, M4 = M8H0.5.

This early guidance translated into accelerated surface colonization. Micron-scale patterns (M1–M2) accelerated surface coverage at days 3–14 across all ELR hydrogels, with ELR2 consistently reaching the highest coverage and shortest time-to-confluence across patterns. By day 3, ELR-based hydrogels, particularly ELR2, supported significantly greater cell coverage than gelatin (**Fig. 5A** and **Fig. 6B**). This divergence widened over time. By day 7, grooved ELR substrates (especially M1-M2) approached confluence, while gelatin lagged significantly (**Fig. 5B** and **Fig. 6B**). By day 14, ELR hydrogels were near-confluent across most patterns, whereas gelatin remained sub-confluent on all but the most potent topographies (M1-M2) (**Fig. 6A,B**). Analysis of cumulative coverage (AUC) and time-to-confluence (T90/T95) confirmed that the combination of ELR chemistry (particularly ELR2) and the 1 µm-high micron-scale topographies (M1–M2) yielded the most rapid and complete monolayer formation (**Fig. 6C** and **Fig. 7**).

**Figure 6.**
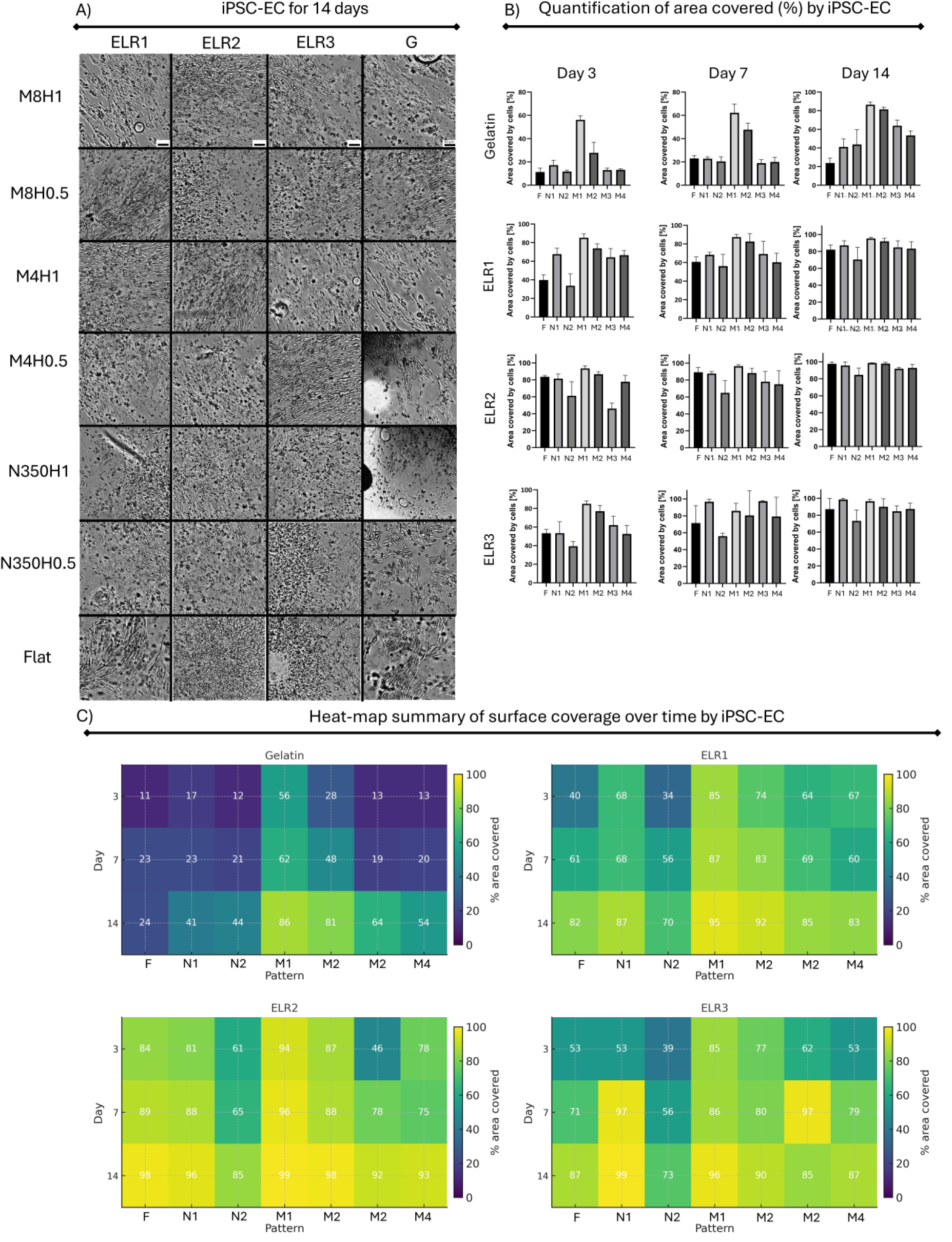
| Patterned ELR-based hydrogels accelerate expansion to confluence relative to gelatin. (A) Representative bright-field mosaics at day 14 for Gelatin vs ELR1–3 across F, N1/N2, M1–M4. (B) Quantitative analysis of surface coverage (% area) at days 3, 7, and 14 (bars = mean ± SD; points = replicates). (C) Heat-map summary of surface coverage over time (Day × Pattern per material; Gelatin, ELR1–3); numbers indicate mean % coverage. Micron grooves (M1–M2) consistently promote faster expansion, with ELR2 achieving the shortest time to near-confluence. Statistics: two-way ANOVA per day (Material × Pattern) with Šídák correction for (a) each pattern vs Flat within a material and (b) each ELR vs Gelatin within a pattern (see legend and SI).

**Figure 7.**
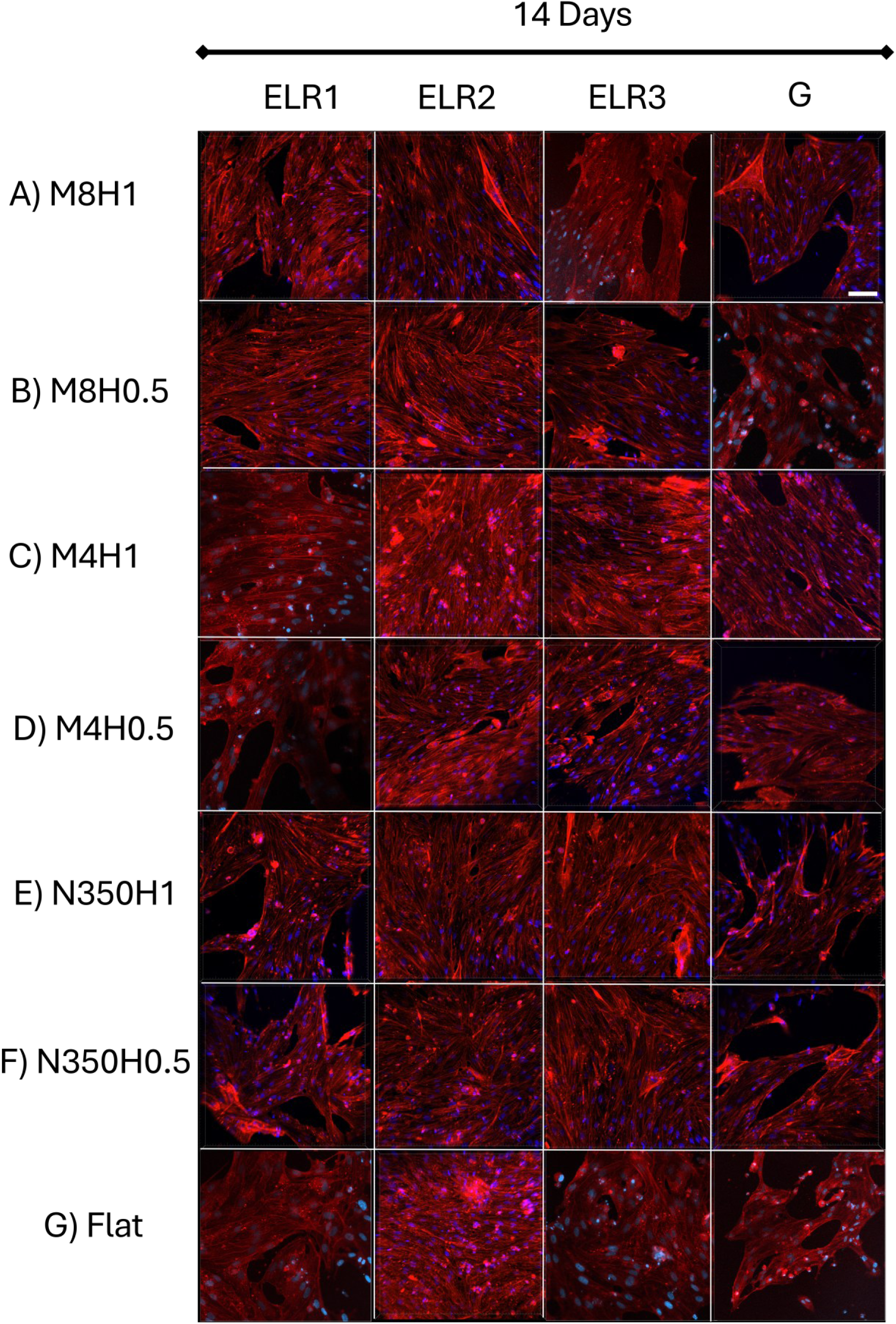
| F-actin organization and contact guidance on patterned ELR-based hydrogels (day 14). (A) Representative immunofluorescence images of iPSC-ECs cultured for 14 days on patterned ELR1, ELR2 and ELR3 hydrogels, compared with gelatin. Patterns include flat (F), sub-micron grooves (N1, N2), and micron-scale grooves (M1–M4). F-actin (phalloidin, red); nuclei (DAPI, blue). Cells on ELR hydrogels form dense monolayers with elongated stress fibers aligned with the groove axis—most pronounced on M1–M2, intermediate on N1–N2, and largely isotropic on flat. Gelatin shows discontinuous coverage with disorganized actin and frequent gaps. Scale bar, 100 µm.

### Cytoskeletal organization in confluent monolayers

After 14 days, F-actin staining revealed nearly continuous monolayers with well-developed stress fibers on ELR hydrogels. On micron-scale grooves (M1–M2) the cytoskeleton was highly organized, with dense, parallel actin stress fibers aligned to the groove axis. Flat ELR substrates yielded confluent cell monolayers with largely isotropic actin organization. In contrast, cells on gelatin were sparse, poorly spread, and exhibited disordered actin networks (**Fig. 7**). These structural observations corroborate the quantitative area-coverage and orientation data (**Fig. 6C and Supplementary Fig. S7–S10)**, confirming that patterned ELR-based hydrogels direct the formation of a stable, aligned endothelium.

## DISCUSSION

This study shows that matrix chemistry and surface topography jointly regulate endothelial cell behavior, influencing outcomes from initial attachment to final confluence. We have demonstrated a materials–geometry regime on soft, protein-mimetic hydrogels in which sequence-defined ELRs and micro/nano-gratings act in concert to dramatically accelerate iPSC-EC capture, alignment, and monolayer formation. The novelty of our approach lies in combining a rapid, minutes-scale imprinting workflow with a tunable hydrogel chemistry, creating a platform that is both optically clear for live imaging and directly portable to curved, lumen-like surfaces [3], [10], [4], [9], [12].

Our stringent 15-minute attachment assay revealed that ELR chemistry provides a critical baseline for cell adhesion, with all ELR coatings significantly outperforming gelatin. This chemistry-driven advantage was further modulated by topography, as specific groove regimes (notably the sub-micron N2 and micron-scale M1 patterns) enhanced early cell retention. This immediate effect is consistent with established contact-guidance models, where topographic features bias protrusion dynamics and stabilize nascent focal adhesions against fluid shear, a phenomenon previously documented on rigid substrates [2], [4], [9] and recently extended to soft hydrogels showing anisotropic traction responses [12], [14], [19]. Crucially, these early advantages were not transient; they propagated to the tissue scale. ELR2 and ELR3 on M1/M2 grooves supported the development of confluent, aligned monolayers by day 14, while gelatin remained sparsely populated under identical conditions. Although ELR3 contains the highest density of RGD motifs, the protease-sensitive ELR2 variant achieved superior endothelial coverage, likely due to a favorable balance between adhesion strength, matrix remodeling, and topographic fidelity [5]–[7], [9], [11], [15], [21]. This demonstrates that the initial biophysical cue is locked in through subsequent proliferation and junction maturation, a process enhanced by groove geometries known to polarize the cytoskeleton and direct collective cell migration [3], [10], [4], [9], [19], consistent with junctional remodeling driven by coordinated actomyosin forces along groove orientation [16].

The superior performance of ELR composites is underpinned by their mechanical and structural properties. Rheology confirmed them as elastic-dominant, shear-thinning networks, capable of storing elastic energy while permitting local remodeling under cell traction. Cryo-SEM revealed a fibrillar, interconnected architecture, and tensile testing delineated the mechanical contributions of each ELR: while ELR3 achieved the highest ultimate strength, ELR2 provided the most consistent and highest small-strain stiffness. This reproducible mechanical integrity, particularly of ELR2, likely resists micro-tearing during critical phases of colony expansion and merger, preventing the adhesion resets that plague weaker gelatin matrices [3], [10], in agreement with prior work on sequence-controlled ELRs where tailored stiffness preserves endothelial integrity under cyclic strain [5]–[7], [21], [25].

Furthermore, we found that chemistry gates the geometric response. NMR confirmed successful ELR incorporation into the composites and allowed us to discriminate between formulations. Functionally, the RGD motifs in ELR3 promote robust initial integrin engagement, whereas the protease-responsive DRIR sequence in ELR2 appears to slow interfacial remodeling, preserving both ligand availability and topographic fidelity during the critical seeding window. Notably, the RGD-bearing ELR3 did not consistently outperform the uPA-responsive ELR2, despite providing a defined integrin-binding motif. This is likely because gelatin is inherently cell-adhesive and already supplies multiple integrin-recognition sequences, making the additional RGD partially redundant in this composite context. In contrast, ELR2 introduces a genuinely new functionality, uPA-mediated proteolytic remodeling, that is not present in gelatin, which may explain its superior performance in supporting endothelial spreading and monolayer formation on selected patterns. Overall, the consistent outperformance of ELR2 on key patterns suggests that controlling the degradation timeline can be as important as maximizing initial adhesion density for ensuring durable alignment and rapid coverage [2], [4], [9], [18], [19].

From a practical standpoint, the identified ELR2/ELR3 × M1/M2 combinations (micro-size grooves of 1 µm) define a readily deployable design window. These pairs are favorable for creating expansion surfaces that minimize pre-culture handling, optically clear organ-on-chip inserts, and pre-aligned luminal liners for vascular grafts—addressing a long-standing gap dominated by opaque, rigid materials [1], [23]. Conceptually, our work demonstrates that temporal coupling from seconds to weeks can be engineered by pairing sequence-encoded matrix properties with feature sizes that match cellular sensing scales [3], [10], [4], [9], linking nanoscale integrin clustering (∼100–350 nm) to multicellular alignment and junction co-orientation on micron-scale patterns [13], [14], [16], [24].

### Limitations and future outlook

All biological experiments in the present study were performed under static culture conditions in order to isolate the effects of matrix bioactivity and surface topography on endothelial attachment and organization. It is well established that endothelial maturation and barrier function are strongly influenced by physiological shear stress. Future studies will therefore investigate the performance of patterned ELR hydrogels under dynamic flow conditions and in curved geometries relevant to vascular constructs. Experiments in our laboratory suggest that the patterned ELR matrices remain mechanically stable under perfusion, supporting the formation of aligned endothelial monolayers. These studies are currently being expanded toward perfusable systems and tubular geometries.

This study, however using a single iPSC-EC line under static conditions, provides a foundational map. The logical next steps involve introducing physiological complexity testing barrier function (TEER, VE-cadherin continuity), nitric oxide release, and adhesion stability under pulsatile flow, especially in scenarios where flow direction is aligned or misaligned with topographical guidance [4], [9], and examining how flow-induced stress couples with substrate anisotropy to regulate VE-cadherin mechanotransduction [12], [16]. Pattern fidelity, particularly for the smallest nanogrooves, could be further optimized with colder imprinting temperatures. Finally, validating this platform across higher aspect ratio designs (perhaps beneficial for cell retention) and multiple iPSC-EC lines will be essential to demonstrate its generality [8]. This workflow is also compatible with gently curved geometries, such as tubular molds, but these demonstrations remain to be validated experimentally in future work.

### Conclusions

This study introduces patterned ELR–gelatin composite hydrogels as a versatile platform for engineering endothelial interfaces. By integrating programmable recombinamer bioactivity with micro- and nanoscale surface architectures, the system enables rapid endothelial cell capture, guided cytoskeletal alignment, and accelerated monolayer formation. The results highlight how matrix bioactivity, mechanics, and topography can be combined to regulate endothelial organization on soft hydrogel substrates. This materials-design strategy provides a foundation for future development of vascular biomaterials and microphysiological systems requiring controlled endothelialization.

## AUTHOR CONTRIBUTIONS

J.L. and A.R., conceptualized and designed the experiments.

Y.R., M.M., K.T., P.P., D.U., and I.P. were involved in the methodology.

J.L., J.C.R.-C., and A.R. provided resources.

J.L. wrote the original draft. All authors were involved in writing, review and editing.

J.L. performed statistical analysis.

J.L. and A.R. acquired funding and supervised the project. All authors agreed to the authorship.

## DATA AVAILABILITY

All data supporting the findings of this study are presented within the manuscript and its supplementary files. Any additional information or inquiries can be directed to the corresponding authors upon reasonable request.

## ACKNOWLEDGEMENTS

The authors are indebted to the European Union’s Horizon 2020 research and innovation programme under the Marie Skłodowska-Curie Actions (Grant Agreement No. 101025242) and the National Science Centre, Poland (Grant No. 2022/47/D/ST8/03467).

This research was partially supported by the Spanish Ministry of Science, Innovation and Universities-MICIU (PID2021-123925OB-I00 and PID2024-15802908OB-I00), AGAUR (2021-SGR-974), Fundació La Marató de TV3 (202331-30), Fundació “La Caixa” (LCF/PR/HR23/52430011), and the CERCA Programme/Generalitat de Catalunya.

JCRC acknowledges support from the Spanish Government (PID2022-137484OB-I00 funded by MCIN/AEI/10.13039/501100011033 and ERDF), the Junta de Castilla y León (VA188P23 and CLU-2023-1-05, co-funded by ERDF), and the Centro en Red de Medicina Regenerativa y Terapia Celular de Castilla y León. PP acknowledges financial support from statutory funds of the Poznań University of Technology, granted by the Polish Ministry of Science and Higher Education (No. 0612/SBAD/3640).

