## Supplementary Figures and Tables for "Patterned ELR-Gelatin Hydrogels Enable Rapid Endothelial Monolayer Formation via Bioactive Matrix Chemistry and Surface Topography"

**Supplementary Table S1 –** Mapping between full pattern identifiers and abbreviated codes

| Abbreviation | Full name | Groove width | Height |
|---|---|---|---|
| F | Flat | — | — |
| N1 | N350H1 | 350 nm | 1 µm |
| N2 | N350H0.5 | 350 nm | 0.5 µm |
| M1 | M4H1 | 4 µm | 1 µm |
| M2 | M8H1 | 8 µm | 1 µm |
| M3 | M4H0.5 | 4 µm | 0.5 µm |
| M4 | M8H0.5 | 8 µm | 0.5 µm |

**Supplementary Table 2**. List of antibodies and primers.

| Antibody | Type/Use | Host | Dilution | Manufacturer |
|---|---|---|---|---|
| VE-cadherin(IgG) | Primary/IF | Rabbit | 1:200 | Invitrogen, PA5-19612 |
| Phalloidin, Texas Red® | -/IF | - | 1:40 | Life technologies, T7471 |
| Donkey anti-rabbit IgG | Secondary/IF | Donkey | 1:200 | Jackson, 711-545-152 |

| Primer | Sense | Antisense |
|---|---|---|
| ***PECAM1*** | AGTCGGACAGTGGGACGTAT | ATGACCTCAAACTGGGCATC |
| ***CDH5*** | GATTTGGAACCAGATGCACA | ACTTGGCATTCTTGCGACTC |
| ***GAPDH*** | GCACCGTCAAGGCTGAGAAC | AGGGATCTCGCTCCTGGAA |

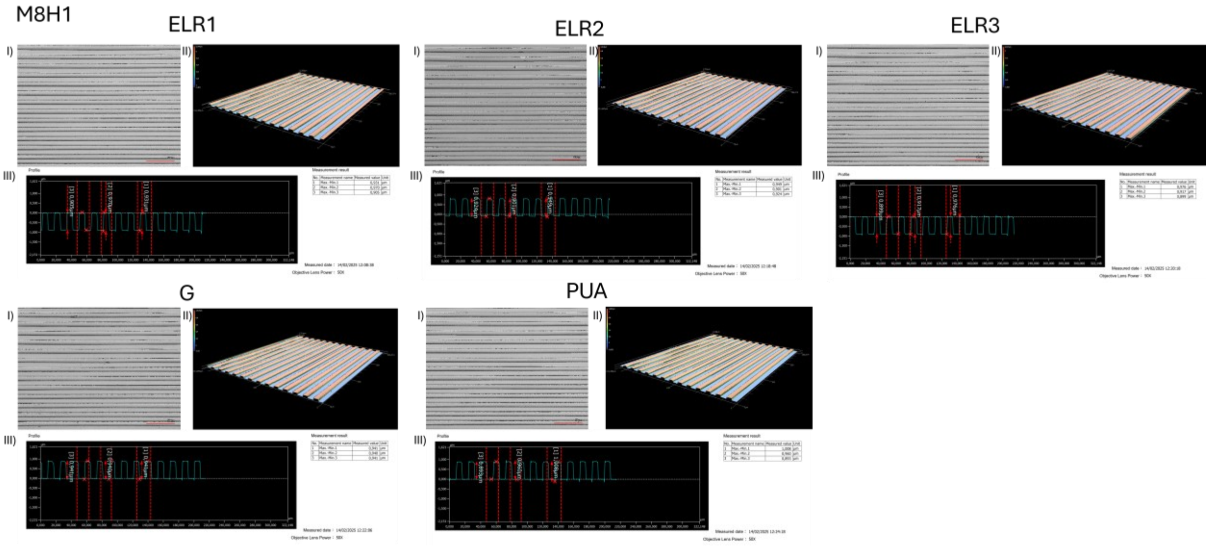


Figure S1 - Surface topography of hydrogel replicas imprinted with line gratings M8H1. Representative bright-field micrographs of ELR1, ELR2, ELR3, gelatin (G), and the PUA master mold, alongside 3D height maps and reconstructions acquired with a digital microscope (VHX-2000, Keyence). Imprints reveal faithful transfer of microscale patterns across different hydrogel formulations.

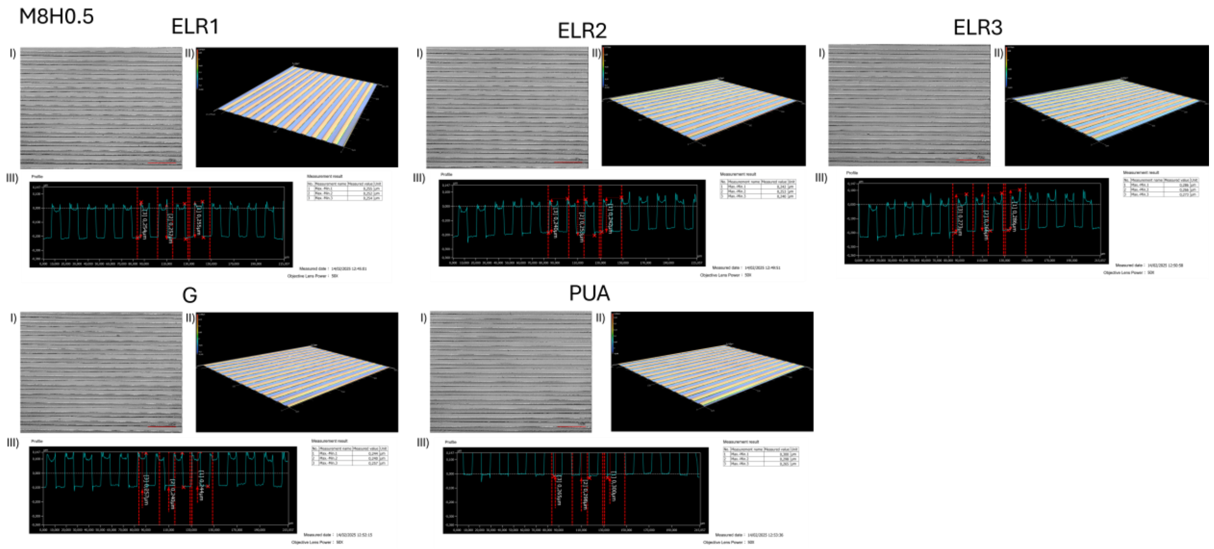


Figure S2 - Surface topography of hydrogel replicas imprinted with line gratings M8H0.5. Representative bright-field micrographs of ELR1, ELR2, ELR3, gelatin (G), and the PUA master mold, alongside 3D height maps and reconstructions acquired with a digital microscope (VHX-2000, Keyence). Imprints reveal faithful transfer of microscale patterns across different hydrogel formulations.

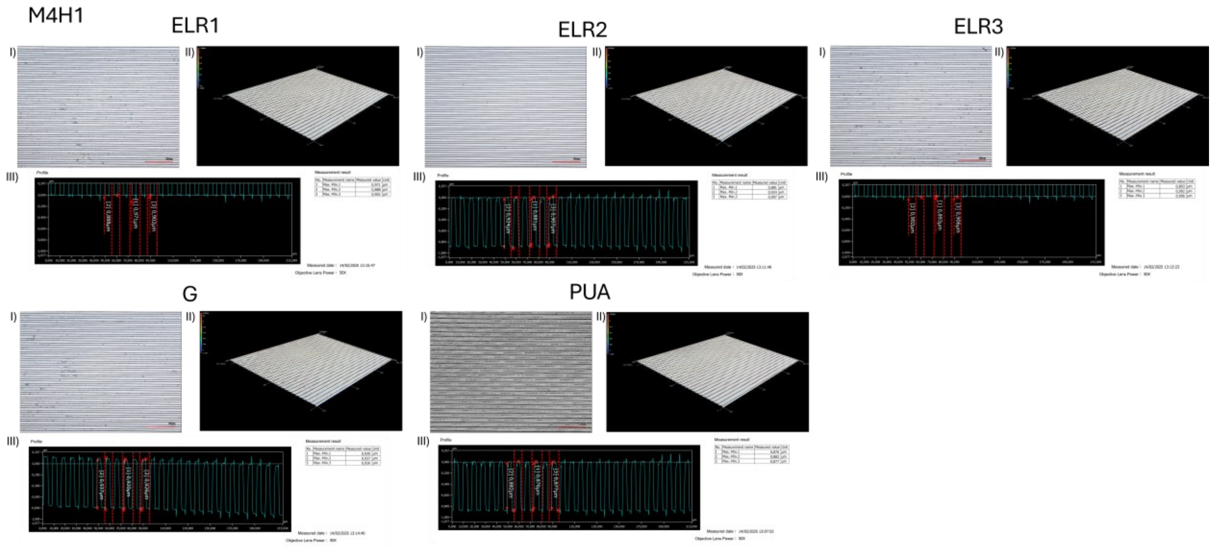


Figure S3 - Surface topography of hydrogel replicas imprinted with line gratings M4H1. Representative bright-field micrographs of ELR1, ELR2, ELR3, gelatin (G), and the PUA master mold, alongside 3D height maps and reconstructions acquired with a digital microscope (VHX-2000, Keyence). Imprints reveal faithful transfer of microscale patterns across different hydrogel formulations.

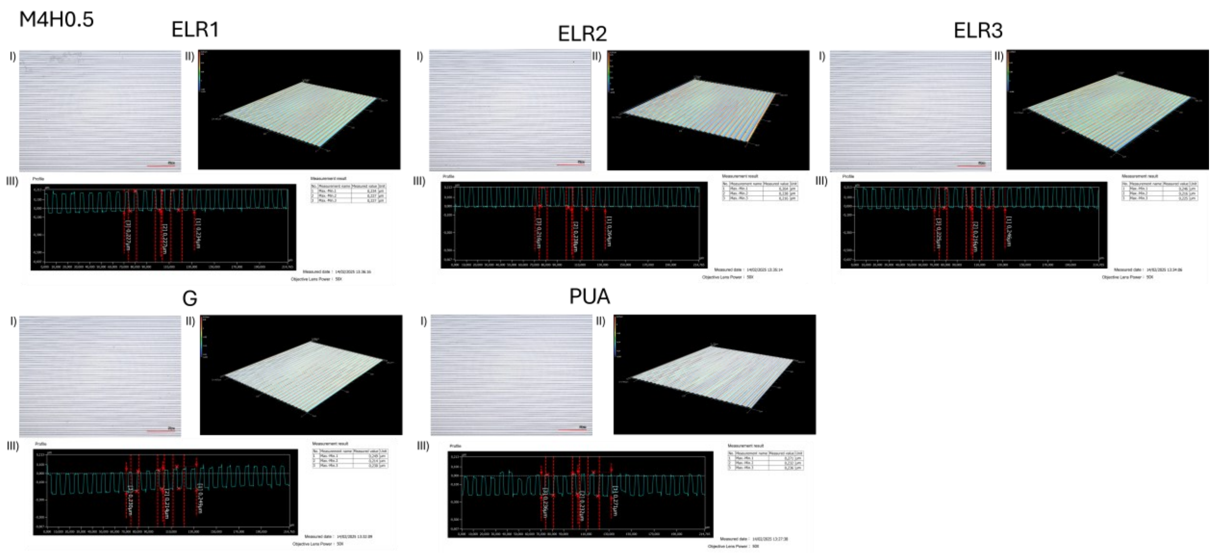


Figure S4 - Surface topography of hydrogel replicas imprinted with line gratings M4H0.5. Representative bright-field micrographs of ELR1, ELR2, ELR3, gelatin (G), and the PUA master mold, alongside 3D height maps and reconstructions acquired with a digital microscope (VHX-2000, Keyence). Imprints reveal faithful transfer of microscale patterns across different hydrogel formulations.

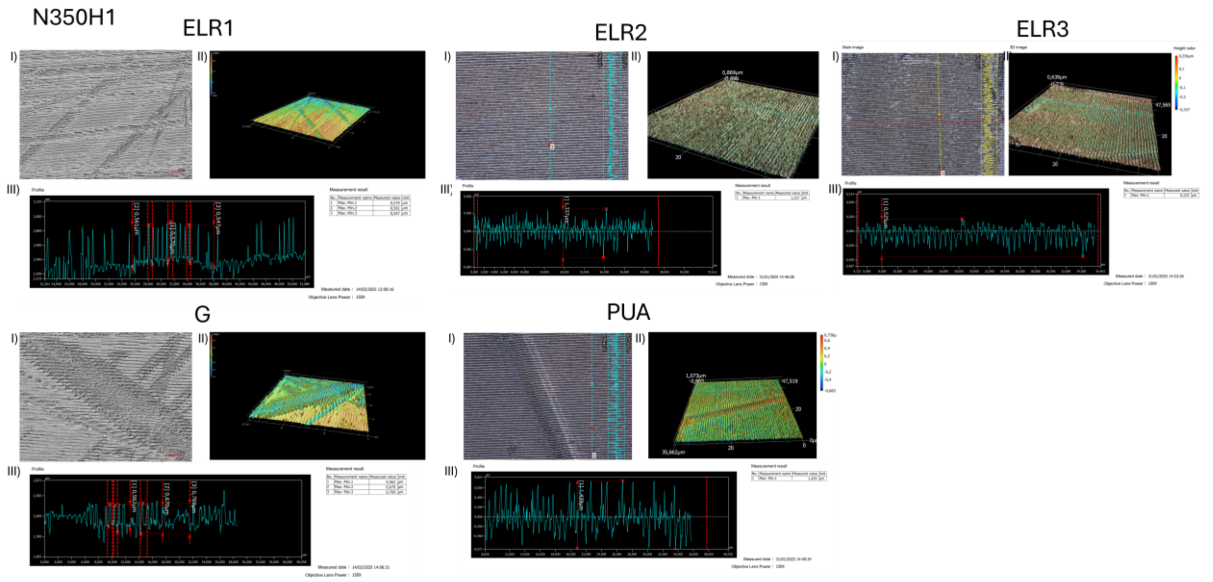


Figure S5 - Surface topography of hydrogel replicas imprinted with line gratings N350H1. Representative bright-field micrographs of ELR1, ELR2, ELR3, gelatin (G), and the PUA master mold, alongside 3D height maps and reconstructions acquired with a digital microscope (VHX-2000, Keyence). Imprints reveal faithful transfer of microscale patterns across different hydrogel formulations.

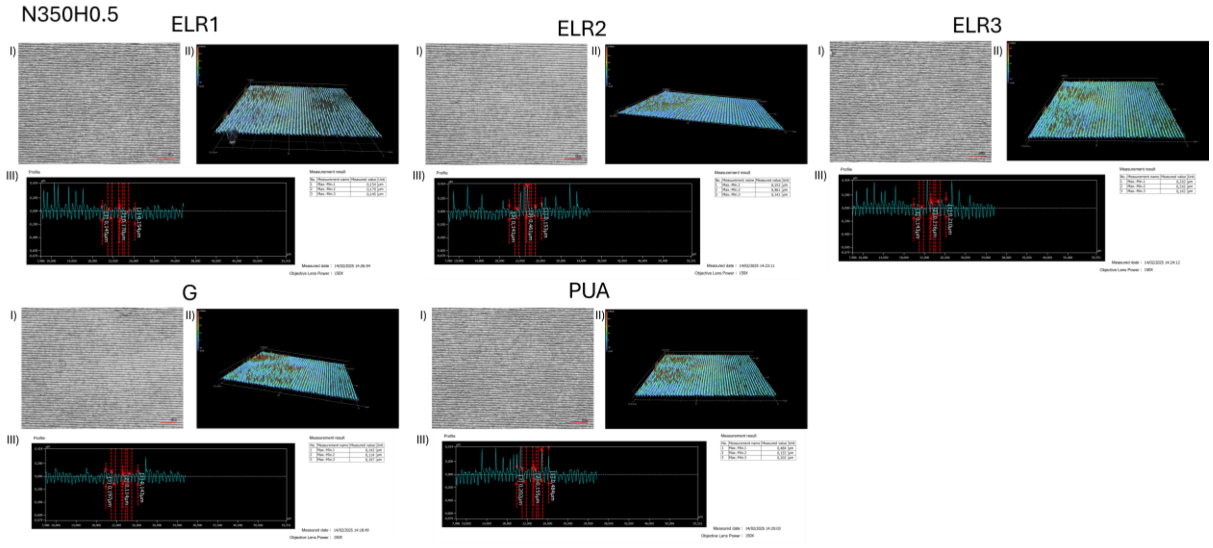


Figure S6 - Surface topography of hydrogel replicas imprinted with line gratings N350H0.5. Representative bright-field micrographs of ELR1, ELR2, ELR3, gelatin (G), and the PUA master mold, alongside 3D height maps and reconstructions acquired with a digital microscope (VHX-2000, Keyence). Imprints reveal faithful transfer of microscale patterns across different hydrogel formulations.

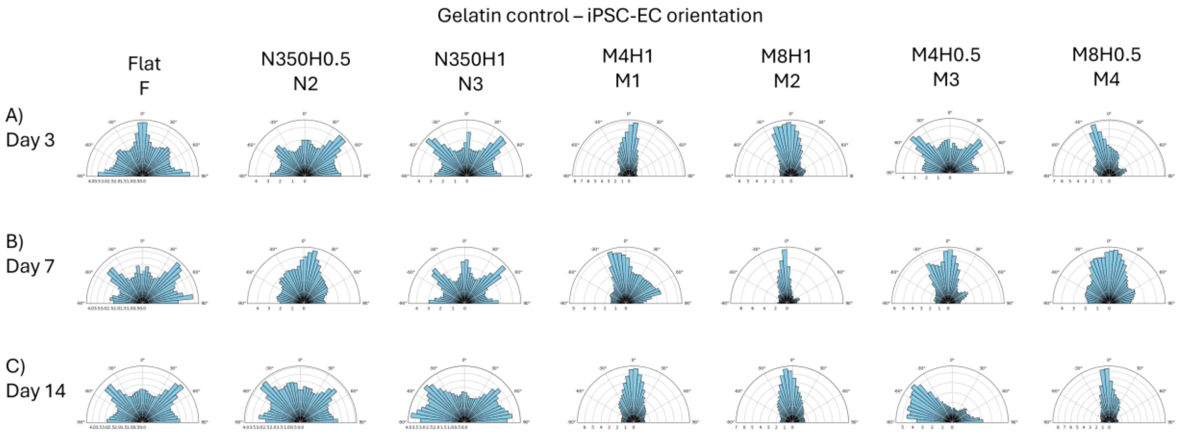


Figure S7 - Circular histograms showing the orientation distribution of iPSC-derived endothelial cells (iPSC-ECs) cultured under the indicated conditions on gelatin control. Each plot represents the frequency of cell alignment angles relative to the pattern direction.

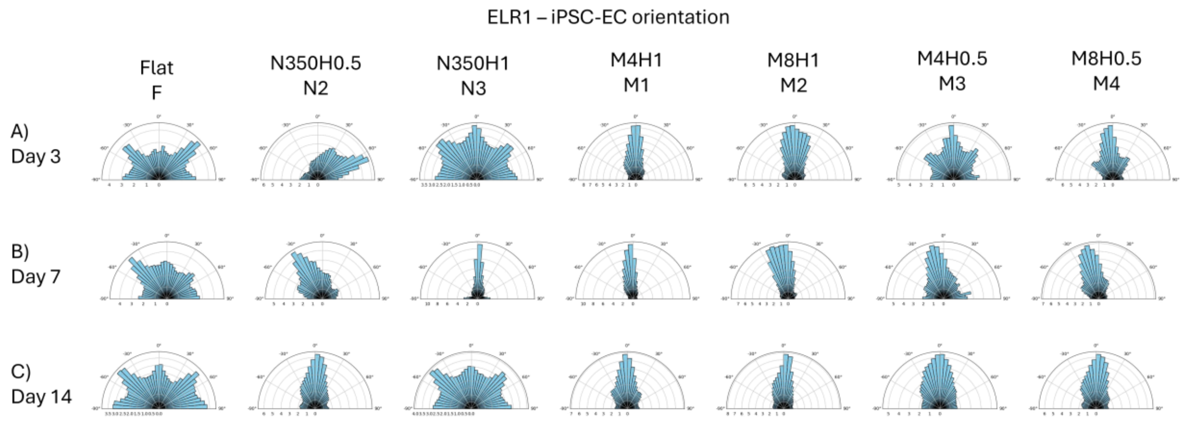


Figure S8 - Circular histograms showing the orientation distribution of iPSC-derived endothelial cells (iPSC-ECs) cultured under the indicated conditions on ELR1 hydrogels. Each plot represents the frequency of cell alignment angles relative to the pattern direction.

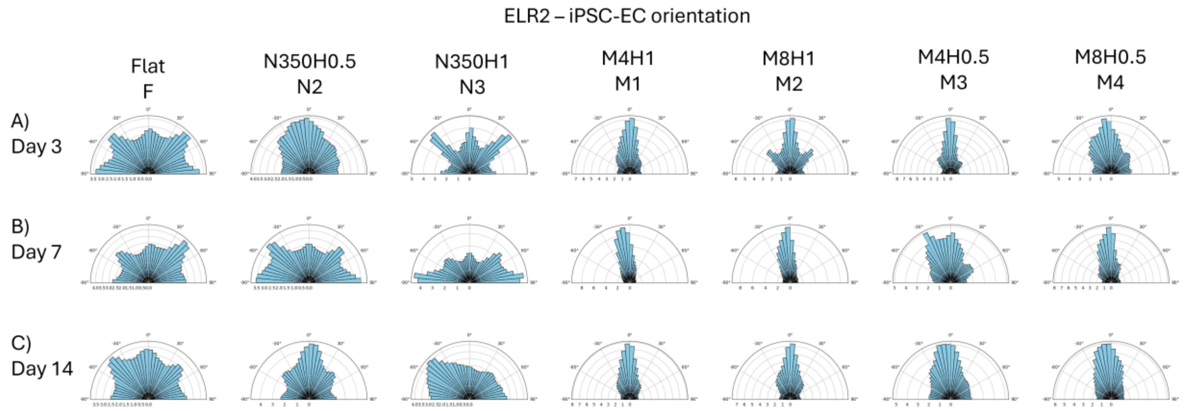


Figure S9 - Circular histograms showing the orientation distribution of iPSC-derived endothelial cells (iPSC-ECs) cultured under the indicated conditions on ELR2 hydrogels. Each plot represents the frequency of cell alignment angles relative to the pattern direction.

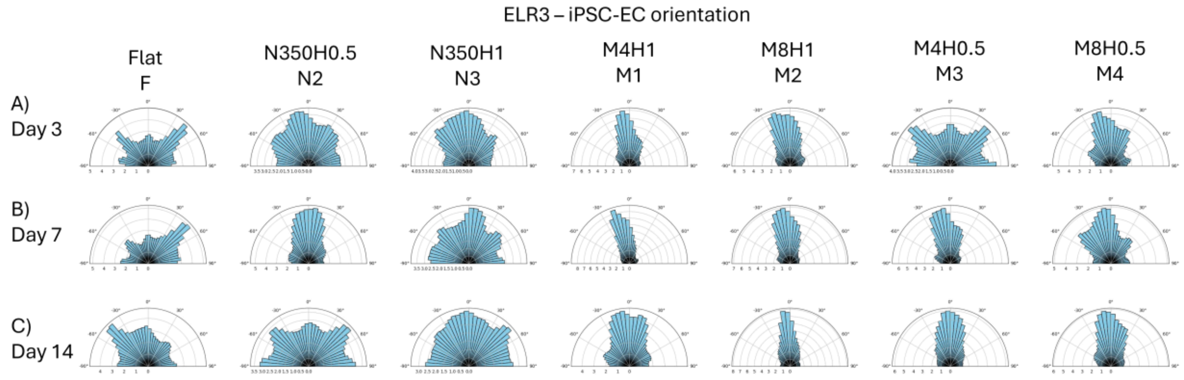


Figure S10 - Circular histograms showing the orientation distribution of iPSC-derived endothelial cells (iPSC-ECs) cultured under the indicated conditions on ELR3 hydrogels. Each plot represents the frequency of cell alignment angles relative to the pattern direction.
